# Assessment of chemical methods in the extraction of spore surface layers in *Clostridioides difficile* spores

**DOI:** 10.1101/2025.08.19.671018

**Authors:** Javier Sanchez, Alba Romero-Rodriguez, Scarlett Troncoso-Cotal, Morgan S. Osborne, Theresa Ariri, Joseph A. Sorg, Daniel Paredes-Sabja

**Author notes:** Corresponding Author. Dr. Daniel Paredes-Sabja, Department of Biology, Texas A&M University, College Station, TX, 77843, USA. Present address: Department of Molecular Biology and Biotechnology, Institute of Biomedical Research National Autonomous University of Mexico (UNAM), City of Mexico. Present Address: Department of Cellular and Molecular Biology, Pontificia Universidad Católica de Chile (Scarlett Troncoso-Cotal).

## Abstract

*Clostridioides difficile* spores are essential for initiation, recurrence, and transmission of *C. difficile* infections (CDI). These outermost layers of the spore, the exosporium and spore coat, are responsible for initial interactions with the host and spore resistance properties respectively. Several spore coat /exosporium extraction methods have been utilized to study the spore surface with differing procedures making comparison across studies difficult. Here, we tested how commonly used exosporium and spore coat extraction methods, termed EBB, USD, and Laemmli, remove the spore coat and exosporium layers of *C. difficile* spores. We assessed the impact of these extraction methods on the spore through transmission electron microscopy, phase contrast microscopy, western blotting, and lysozyme triggered cortex degradation. Transmission electron microscopy shows that treatment with EBB and USD, completely remove the spore coat and exosporium layer while leaving decoated spores intact. Western blots revealed differences in the ability to extract spore surface protein markers (CdeC, CdeM, CotA). In addition, lysozyme was able to degrade the cortex in decoated spores regardless of the treatment employed. Western blot analysis of lysozyme treated-decoated spores, reveals that EBB and USD treatment allows for detection and release of the spore core germination protease, GPR. Our results provide a comparison of commonly used extraction methods in *C. difficile* spore biology, standardizing their impact in spore coat and exosporium extraction for use in future studies.

**Importance:** The outermost layers of *C. difficile* spores, the exosporium and spore coat, are essential for the spores resistance properties and initial interactions with the host. However, there is variability in extraction protocols making it difficult to compare across studies. This work evaluates the commonly used extraction methods EBB, USD, and Laemmli at removing the exosporium and spore coat and provides a foundation for improved reproducibility. Here, we identified the effectiveness of these different extraction methods allowing us to better understand these techniques to accurately analyze spore surface in *C. difficile* spore research.

## Introduction

*Clostridioides difficile* is a Gram-positive anaerobe that has become a leading cause of antibiotic associated diarrhea in developed countries^1, 2^. Treatment against *C. difficile* infections (CDI) resolve 95% of primary cases; however, ∼15 – 30% of recovered CDI patients develop subsequent recurrent episodes of CDI, which result in mortality rates increasing up to 30%^3^. Although two major clostridial toxins, TcdA and TcdB, are necessary for disease manifestation^4^, the production of *C. difficile* spores during infection is essential for disease recurrence^5, 6^.

*C. difficile* spores are metabolically dormant and naturally resistant to antibiotics, heat, ethanol, and various cleaning agents^6, 7^. The spores are made up of a series of concentric layers that assemble under the control of RNA polymerase sporulation-specific sigma factors, SigF, SigE, SigK, and SigG ^6, 7^. Importantly, during late stages of sporulation, *C. difficile* forms two distinctive spores that differ in the thickness of the exosporium morphotype, while the underlying spore coat remains identical ^6, 8, 9^.The outermost exosporium layer of *C. difficile* spores play important roles in infection, recurrence, and interaction with host molecules ^10–14^. Both, the spore coat and exosporium of *C. difficile* spores are proteinaceous layers made of > 50 and ∼ 10 - 20 proteins, respectively^15, 16^. Both contribute to the spores resistance against various environmental insults^7^. Numerous studies on the spore coat and exosporium layer of *C. difficile* spores have varied in the methods to extract the spore coat and exosporium layers ^12, 15–23^. Because these methods dissociate the entire proteinaceous spore surface layers, they are thought to extract the spore coat and exosporium layers, at least in *C. difficile*^12, 15–23^. Although different in their components, most of these chemical approaches use: i) a chaotropic agent, urea or urea + thiourea to disrupt hydrogen bonds and results in the dissolution of hydrophobic residues within the protein^24^; ii) SDS to denature and solubilize proteins; iii) a reducing agent which can be 2-mercaptoethanol or dithiothreitol. Currently, three methods have been commonly utilized to remove the spore surface layers in *C. difficile* spores. Several studies utilize 2 X Laemmli buffer (1% SDS, 5% 2-mercaptoethanol) followed by boiling for 10 min at 100°C ^12, 15, 17^ whereas other proteomic studies have used lysis buffer with 8 M urea, and DTT as a reducing agent^16, 21, 22^. In *C. difficile* spore research, of 8 M urea, 2 M thiourea, 5% (w/v) SDS, 2% 2-mercaptoethanol, denominated EBB buffer, followed by incubation for 10 min at 95 °C has also been used extensively for extraction and solubilization of spore proteins mainly in sporulating cultures ^18–20^. In addition, urea-SDS-DTT denoted as USD (8 M Urea, 50 mM DTT, 1% w/v SDS) has been used in the extraction of exosporium and spore coat in C. difficile spores^25^. Despite widespread use of these different chemical extraction methods, it is unclear whether these chemical methods, commonly employed in *C. difficile* spore-research, efficiently remove the spore coat and exosporium layer.

Efficient fractionation of the spore surface layers is essential to understand the localization of spore surface constituents in each of these layers. In this regard, we demonstrated that enzymatic digestion with proteinase K or trypsin, or a mechanical treatment with sonication, efficiently removed only the exosporium layer of *C. difficile* spores ^15^. This allowed the identification of the exosporium proteome in spores of strain 630^15^; however, a similar validation with chemical extraction methods has not been conducted. Therefore, to improve our tool-box of methods to study *C. difficile* spore architecture and composition, in this work, we tested commonly used chemical extraction methods, EBB, USD, and Laemmli in their ability to remove the spore coat and exosporium layers and their impact on spore structure. By utilizing antibodies against exosporium markers (i.e, CdeC and CdeM)^12, 26, 27^, spore coat specific marker CotA, the spore cortex lytic enzyme SleC, and germination protease GPR as a core specific marker^28, 29,30^ we evaluated how these commonly utilized chemical methods to extracted these layers through immunoblotting, transmission electron microscopy (TEM), and lysozyme permeability. For comparison we defined “extraction efficiency” as a measure of ability extract protein as seen through SDS-PAGE and immunoblotting, complete removal of exosporium and spore coat as seen through TEM, removal of spore surface markers (CdeM, CdeC, CotA, SleC, and GPR) from decoated spore pellets and lastly removal of spore coat following lysozyme treatment.

## Materials and Methods

### Bacterial strains and growth conditions

Bacterial strains used in this study are listed in Main Table 1. *C. difficile* R20291 strain was grown at 37°C under anaerobic conditions (Coy Laboratory anaerobic chamber, 4% H_2,_ 5% CO_2_, 85% N_2_) on Brain Heart Infusion supplemented with 0.5% yeast extract, and 0.1% L-cysteine (BHIS) broth or 1.5% agar. *E. coli* DH5α strains were grown on Luria Bertani supplemented with 50 ug/mL chloramphenicol, 10 ug/mL tetracycline, 50 ug/mL kanamycin, or 100 ug/mL ampicillin where indicated.

**TABLE 1.**
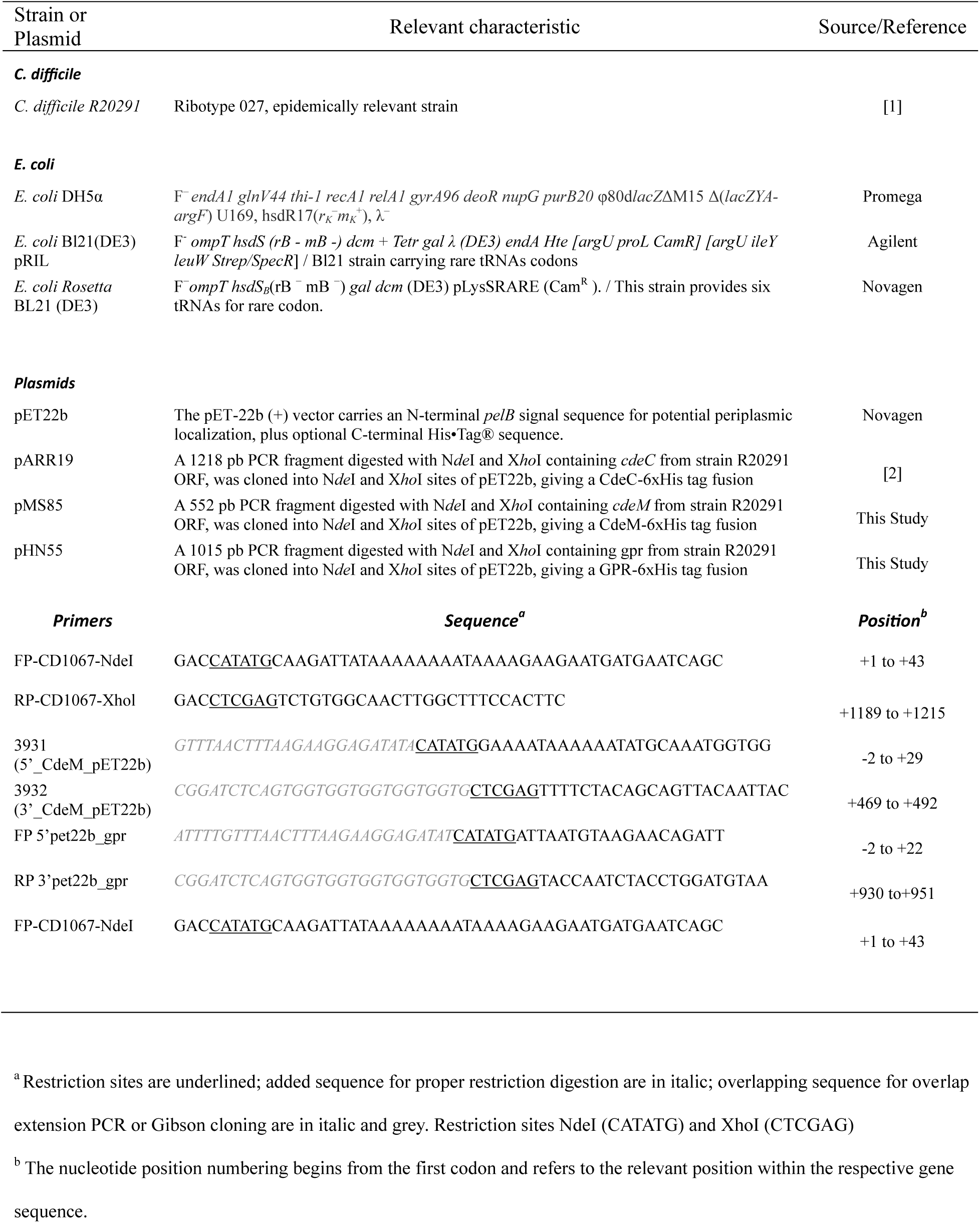
Bacterial strains and plasmids used.

### Plasmid construction

The plasmids pARR19, pMS85, and pHN55 were made from pET22b as follows and listed in Main Table 1. pET22b was digested with NotI / XhoI and purified by gel electrophoresis, extracted, then assembled through restriction cloning with *cdeM* or *cdeC,* or *gpr* fragment via Gibson assembly. The oligonucleotide primers 3931 (5’_CdeM_pET22b) and 3932 (3’_CdeM_pET22b) were used to amplify *cdeM*,primers FP-CD1067-NdeI and RP-CD1067-XhoI were used to amplify *cdeC,* and (FP-5’pet22b_gpr and RP RP 3’pet22b_gpr were used to amplify *gpr* from *C. difficile* R20291. After Gibson assembly and transformation into *E. coli* DH5α, colonies were re-streaked and tested via PCR for correct assembly, followed by whole plasmid sequencing using Oxford Nanopore Technology by Plasmidsaurus Inc. (Eugene, OR).

### Spore preparation and purification

*C. difficile* strain R20291 was grown in Brain Heart Infusion broth (BHIS) (Difco) supplemented with 0.5% yeast extract and 0.1% cysteine anaerobically at 37°C. Overnight cultures were grown in BHIS broth with 0.1% sodium taurocholate and 0.2% d-fructose. Cultures were diluted to an OD_600_ of 0.5 and 250 uL was plated on 70:30 sporulation media under anaerobic conditions for 5 days at 37°C^31^. Sporulating cultures were collected from plates, resuspended in ice-cold MilliQ dH_2_O water, and left in 4°C overnight to allow for lysis of vegetative cells. After incubation, samples were centrifuged, and pellets washed with ice-cold MilliQ dH_2_O water 5 times followed by repeated centrifugation and resuspension to remove vegetative cell debris. Spores were separated using a 60% sucrose gradient and centrifuged for 20 min at 18,400 g, spores were then washed with ice-cold water to remove residual sucrose. To assess purity, spores were imaged on a 1% agarose pad and visualized through phase contrast microscopy. Spores were considered pure once the sample was free of vegetative cells and debris. 5×10^9^ spores/mL of purified spores were quantified using a NeuBauer chamber and stored at-80°C until use.

### Spore extraction methods

1 mL of 5×10^9^ spores/mL were pelleted down at 18,400 g for 5 minutes and resuspended in 500 uL of buffer. Spores with EBB (EBB 8M Urea, 2M Thiourea, 4% w/v SDS, 2% v/v β-mercaptoethanol) were boiled 10 minutes and spun down at 18,400 g for 5 minutes^23^. Spores treated with USD (8M Urea, 1% w/v SDS, 50 mM DTT, 50 mM Tris-HCl pH 8) incubated for 90 minutes at 37°C and pelleted down^32^. All samples were centrifuged using Eppendorf 5425R centrifuge in 1.5mL centrifuge tubes. Supernatant containing spore extract (exosporium, spore coat, cortex fraction) was saved and spore pellet (cortex, core fraction) washed three times with MilliQ dH_2_O water to remove residual spore extract and buffer. Spores were then retreated with EBB for a total of three extractions, with supernatant saved between each step. Laemmli treated spores were boiled for 10 min in 2x Laemmli buffer with 5% (v/v) B-mercaptoethanol. Supernatant was saved and decoated spore pellet washed three times with MilliQ dH_2_O. Decoated spores from USD and EBB treated three times were incubated with 2mg/ml lysozyme at 37°C for 2 hours to degrade cortex and core for immunoblotting^30^.

### Germination assay and lysozyme resistance

*C. difficile* spores were treated with lysozyme to analyze the effects of the extraction methods on the integrity of the spore coat^31^. Spores were treated once with USD, EBB, or Laemmli and decoated (1×10^7^ spores). Decoated spores were treated with 250 ug/mL lysozyme in 25 mM phosphate buffer (pH 7.4) at 37°C for 30 min and 2 hours^12^. Samples were washed with MilliQ dH_2_O and analyzed under phase contrast microscopy. ∼500 spores were analyzed for germination and binned for phase bright, phase grey, or phase dark at the 2 hour time point. The data represents the results of three independent experiments and spore sample preparations.

### Transmission electron microscopy and analysis

For transmission electron microscopy, we followed prior protocols with modifications^33^. Briefly, *C. difficile* spores were fixed overnight with 3% glutaraldehyde, 0.1 M cacodylate buffer (pH 7.2) at 4°C. Fixed spores were centrifuged at 18,400 g for 5 minutes, supernatant discarded, and stained with 1% osmium tetroxide in 0.05M HEPES buffer (pH 7.4) overnight at 4°C. Treated samples were washed 5x with MilliQ dH_2_O water. The spores were dehydrated stepwise in 30%, 50%, 70%, 90% acetone for 15 minutes respectively, followed by dehydration with 100% acetone 3 times for 30 minutes at each step. A small amount of acetone is left covering the sample to prevent rehydration. Spore samples were embedded in modified Spurr’s resin (Quetol ERL 4221 resin; EMS; RT 14300) in a Pelco Biowave processor (Ted Pella, Inc.). Initially, 1:1 acetone-resin for 10 min at 200W-with no vacuum, 1:1 acetone-resin for 5 min at 200W-vacuum 20”Hg, followed by 100% resin 4 times at 200W for 5 minutes-vacuum 20”Hg. The resin was removed, and sample pellet was transferred to BEEM conical-tip capsule and filled with 100% fresh modified Spurr’s resin. Sample was left to reach bottom of capsule and subsequently left to polymerize for 48 hours in a 65°C oven followed by 24 hours at room temperature. Ultrathin section ∼100nm (silver-gold color) were obtained using a Leica UC7 Ultramicrotome and placed on glow-discharged carbon coated 300-mesh Cu grids. Grids were double lead stained with 2% uranyl acetate for 5 minutes and washed with filter sterilized (0.2uM filter) MilliQ dH_2_O water followed by 5 minute staining with Reynold’s lead citrate and subsequent washing as described. Grids were stored in a dessicator containing phosphorous pentaoxide until ready for imaging. All ultrathin TEM sections were imaged on a JEOL 1200 EX TEM (JEOL, Ltd.) at 100 kV, and images were recorded on an SIA-15C charge-coupled device (CCD) (Scientific Instruments and Applications) camera at the resolution of 2,721 by 3,233 pixels using MaxImDL software (Diffraction Limited). All equipment used is located at the Texas A&M University Microscopy and Imaging Center Core Facility (RRID: SCR_022128).

ImageJ was used to measure all samples in nanometers 10 spores were analyzed for wild type spores and EBB, USD, and Laemmli decoated spores for a total of 40 spores. Initially, the spore and core diameter were measured from cross sections of circular spores three times and the average was calculated. The spore diameter was measured using a straight line through the spore center from one end of the outermost layer to the other. Cortex width was measured once from inner membrane to cortex. Statistical analysis was done using one way analysis of variance (ANOVA) with Tukey test for multiple comparison and ROUT was used to remove outliers. Statistical cutoff for significance with a *P*-value < 0.05 was used.

### Antibody preparation

*E. coli* BL21 (DE3) pRIL was used to overexpress CdeC and CdeM proteins. CdeC and CdeM were respectively overexpressed using pETM11 vector with a C-terminal 6xHIS tag^26^. Plasmids were transformed into BL21 (DE3) pRIL cells and inoculated in 5 mL of Luria Bertani (LB) broth supplemented with 50 ug/mL chloramphenicol, 10 ug/mL tetracycline, and 50 ug/mL kanamycin in a shaker overnight at 37°C. Overnight culture was used to inoculate a 300 mL flask with the necessary antibiotics at a 1:100 ratio and grown at 37°C until OD_600_ reached 0.7-0.8. Fresh LB with the appropriate antibiotics was added until flask reached 1L volume. Once OD_600_ was reached culture was induced with 0.5 mM isopropyl-β-D-1-thiogalactopyranoside (IPTG) for 18 hrs at 21°C. Cultures were pelleted down at 3,968 g for 10 minutes and stored at-80°C. Pellet was resuspended in 5mL of soluble buffer (50 mM NaH_2_PO_4_, 300 mM NaCl, 20 mM imidazole, 1mM phenylmethanesulfonyl fluoride (PMSF), pH 8) and sonicated 15 seconds on, 15 seconds on ice for a total of 6 cycles at 50% amplitude. Lysate was pelleted down at 3,968 g at 4°C for 30 minutes and subsequently filtered with a 0.22 uM filter. Filtered lysate was taken further purified to obtain soluble protein.

Purification of soluble recombinant CdeC and CdeM proteins was done using Akta Start, Cytiva FPLC protein purification system. Briefly, filtered soluble protein extract were loaded on HisTrap FF crude column (GE Healthcare), washed with 15 column volumes (CV) of wash buffer (50 mM NaH_2_PO_4_, 300 mM NaCl, and 20 mM imidazole, pH 8). Proteins were eluted with 10 CV of elution buffer (50 mM NaH_2_PO_4_, 300 mM NaCl, and 250 mM imidazole, pH 8). Samples were eluted as a single peak and subsequent fractions were analyzed by 15% SDS-PAGE and stained with Coomassie G250 to determine purity of fractions. Purified soluble protein was quantified using To Pierce™ BCA Protein Assay Kit (ThermoFisher Scientific). The purified soluble CdeM and CdeC proteins were subsequently used for antibody production. SDS-PAGE and western blot of purified soluble protein provided in Figure S2.

Five BALB/c mice, 7-9 weeks old were used for immunization with purified soluble CdeC and CdeM proteins respectively. A pre-immunized bleed was taken at day-1 from isoflurane anesthetized mice. Blood was incubated for 30 minutes at room temperature to coagulate and then centrifuged at 2348 g for 10 minutes at 4°C. Serum was collected and stored at-20°C. At day 0, 20 ug of recombinant protein was emulsified with Freund’s Complete Adjuvant (FCA) in a 1:1 ratio. Mice were anaesthetized and injected with 200 uL of protein emulsified in FCA subcutaneously in the nape. Mice were inoculated as described at day 14 and 28 however using Freund’s Incomplete Adjuvant (FIA). At day 42, mice were anaesthetized with 5% isoflurane and maintained at 1.5% and a cardiac puncture was performed to collect blood. Serum was collected as previously described. Animal protocol was approved by the Comité de Bioética of the Facultad de Ciencias Biológicas at the Universidad Andrés Bello under the approval act code 0035/2018.

### CdeC, CdeM and GPR expression and purification of inclusion bodies

Plasmids pMS85 and pARR19 were transformed independently into *E. coli Rosetta* BL21 (DE3) and incubated overnight at 37°C on LB-agar supplemented with chloramphenicol (CM, 20 µg / mL) and ampicillin (AMP, 100 µg / mL). A single colony was inoculated into 7 mL of LB broth supplemented with chloramphenicol, ampicillin, and 0.5% glucose. Cultures were grown overnight at 37°C shaking. 100 mL of LB supplemented with CM, AMP, and 0.5% glucose was inoculated with 1 mL of overnight culture. Cultures were grown to an OD_600_ between 0.7-0.9 shaking. Cultures were induced with 0.5 mM IPTG for 4 hours at 37 °C shaking. Cells were pelleted and store at-80°C until purification of the inclusion bodies.

Inclusion bodies were purified as follows. Cultures thawed and resuspended in 15 mL of 10 mM EDTA, 0.1% Tween-20, 1 mM PMSF, and 10 mg/mL lysozyme in PBS and incubated for 1 hr at 37°C. Samples were sonicated at 20% amplitude for 15 seconds on and 3 minutes off 6 times. Pellets were centrifuged at 1,610 x g at 4°C for 10 minutes. Supernatant was discarded and pellets were resuspended in 10 mL of cold PBS with 2% Triton X-100, and 1 mM PMSF. Pellets were centrifuged at 1,610 x g for 10 min at 4°C. Pellets were washed 5 more times. Pellets were resuspended in 2 mL of PBS supplemented with 2% Triton X-100, and 1 mM PMSF. Pellets were sonicated at 20% amplitude 30 seconds on and 30 seconds off for 5 min. 200 µL of sample were pipetted onto 600 µL of 45% histodenz and centrifuged at 5,900 x g for 20 minutes at 4 °C. This was repeated until all debris was removed. Purity of the inclusion bodies was checked under the microscope using phase contrast. Pellets were washed in cold PBS five times and stored at 4°C.

3 mg of sample in PBS supplemented with 10% glycerol were sent to Pacific Immunology for antibody production in rabbits.

Plasmid pHN55 was GPR was into *E.coli Rosetta* BL21 (DE3) on LB-agar supplemented with chloramphenicol and ampicillin. Transformed cells were scraped into 1 mL of LB, back diluted to OD_600_ of 0.01 and used to inoculate 2XTY medium supplemented with ampicillin and chloramphenicol. The culture was incubated at 37 °C with shaking at 190 rpm. Once OD_600_ of the culture reached 0.6 and 0.8, the culture was induced with 1 mM IPTG and incubated, with shaking 180 rpm, for 16 hours at 16 °C.

After induction, the culture was centrifuged for 30 minutes at 6,370 x g and 4 °C. Supernatant was discarded and pellets were stored overnight at-80 °C. One liter of frozen cells was thawed and suspended in 25 mL of LIB buffer (300mM NaCl, 50Mm Tris-HCl, 10% glycerol without imidazole at pH 7.5). After suspension, lysozyme and DNase I was added to each 25 mL of cells and rocked for 30 minutes at 4 °C. Cells were sonicated on ice for 20 minutes at 27% amplitude and centrifuged at 30,285 x g for 30 minutes at 4 °C. The clarified supernatant was added to Ni-NTA agarose beads and incubated overnight with shaking at 4 °C. The beads were washed twice for 15 minutes with LIB buffer supplemented with 15 mM imidazole (pH 7.5) with shaking. The beads were washed once in 1 mL of LIB buffer with 75 mM imidazole (pH 7.5) with shaking for 15 minutes at 4 °C. Beads were eluted with same buffer supplemented with 500 mM imidazole.

Samples were concentrated using 30 kDA molecular weight cut off centrifugal device. 8 mg / ml of expressed protein was aliquoted, and flash frozen in dry ice-ethanol bath, and stored at-80 °C. 3mg of purified protein was sent to Pacific Immunology for generating polyclonal antibody in rabbit. Serum was aliquoted and stored at-80 °C.

### Western blotting

Protein samples were resuspended in 2x SDS-PAGE loading buffer (BioRad) with 5% b-mercaptoethanol and boiled for 10 minutes. Samples were run in a 12% acrylamide SDS-PAGE gels. Proteins were transferred to nitrocellulose membrane and blocked at 4°C overnight in 4% bovine serum albumin (BSA) in Tris-buffer saline (TBS) (pH7.4) with 0.1% TWEEN20 (TBS-T). Western blots probed with primary 1:3,000 anti-CdeC, 1:3000 anti-CdeM, 1:1,000 anti-CotA 1:10,000 anti-SleC^33^, and 1:10,000 anti-GPR produced in rabbit in 1% BSA in TBS-T for 1h at room temperature. SleC was a gift from Dr. Joseph Sorg at Texas A&M University^34^. Membrane was washed three times for 5 minutes with TBS-T and incubated in secondary antibody 1:10,000 dilution goat anti-rabbit HRP (Sigma) in 1% BSA in TBS-T. Membrane was washed three times as described and imaged Licor C-DiGit Blot Scanner.

## Results and discussions

### Experimental design of spore coat and exosporium extraction

Two chemical methods to extract spore coat and exosporium proteins have been extensively utilized in *C. difficile* spore-research^12, 15–23^. However, the extraction efficiency and effect on oligomerization state of spore coat and exosporium proteins using EBB and USD remains unclear. To investigate and compare extraction treatments, we developed an experimental design that involved multiple rounds of extractions, analysis of extracts, as well as decoated spores (Fig 1). As shown in Figure 1, spores were initially treated with either EBB, USD or Laemmli. For EBB treatment, spores were boiled for 10 minutes, centrifuged, and the supernatant containing spore coat and exosporium extracts was saved (1^st^ extraction). A second and third extraction were subsequently performed on the remaining pellets to assess remaining proteins at each stage. For USD treatments, spores were incubated for 90 minutes at 37 °C, whereas for EBB and Laemmli treatments, spores were boiled for 10 min at 100 °C (Fig 1). An aliquot was saved from each step for immunoblotting to determine presence of residual protein markers on the spore surface. The remaining decoated pellet was washed to remove residual buffer and a small aliquot was saved between each treatment (1^st^, 2^nd^, 3^rd^ decoated spore). To analyze proteins within the core and cortex, spores treated three times with EBB or USD were incubated with 2 mg/mL of lysozyme for 2 hours at 37 °C. This design provides a method to interrogate proteins within the spore core and cortex peptidoglycan (PG) layer that were not able to be extracted with EBB, USD, or Laemmli alone. For all treatments, exosporium/spore coat/cortex extract and lysozyme treated decoated spores (cortex/core) fractions were run through SDS PAGE and immunoblotted against specific spore markers (CdeM, CdeC, CotA, SleC, and GPR) (Fig. 1). Moreover, decoated spores were also subjected to TEM and lysozyme-triggered germination for spore core analyses.

**Figure 1.**
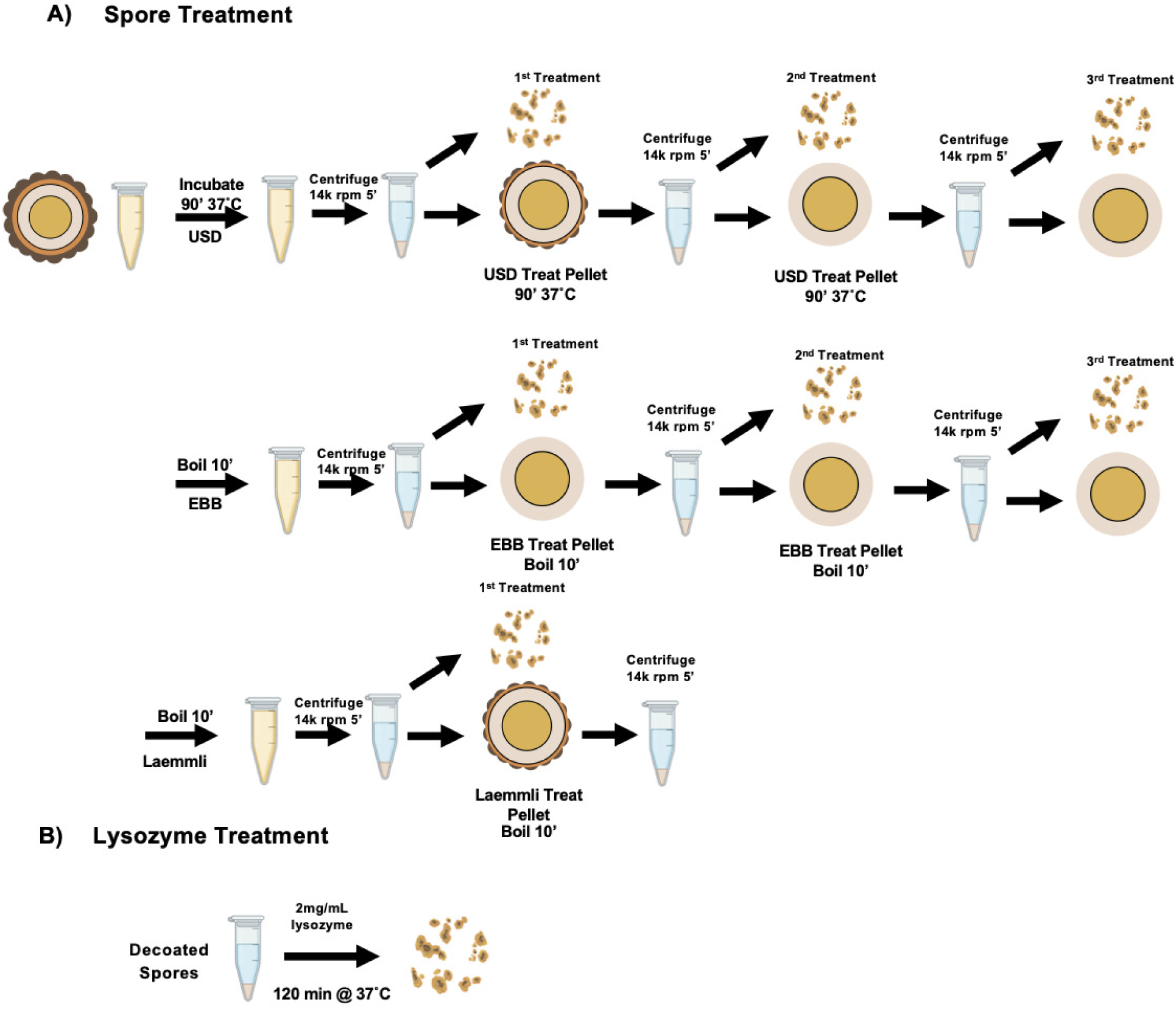
Schematic of extraction of *C. difficile* spores. A) Purified R20291 C. difficile spores (5×10^9^ spores/mL) were pelleted and resuspended in EBB, USD, or Laemmli. EBB and Laemmli treated spores were boiled for 10 minutes while USD treated spores were incubated for 90 minutes at 37°C. After treatment, spores were centrifuged for 5 minutes at 18,400 rcf. The 1^st^ extraction supernatant containing exosporium/spore coat/cortex extracts was collected for SDS-PAGE and Western blotting. The remaining decoated spore pellet was washed three times with sterile MilliQ water and centrifuged at 18,400 rcf for 5 minutes in between washes. The decoated spores were treated to two additional rounds of treatment (2^nd^ and 3^rd^) following the same procedure described above. At each step a portion of the extract and spore pellet were saved for analysis. B) Decoated spores of the 3^rd^ treatment of USD and EBB and 1^st^ Laemmli treatment were incubated with 250 ug/mL lysozyme for 2 hours to asses integrity of the spore coat following treatment.

### Transmission electron microscopy of treated spores and measurements

To examine the effect of extraction on the spore surface ultrastructure, spores treated with a single extraction with EBB, USD, or Laemmli and subsequently decoated were imaged through TEM. As expected, untreated R20291 spores have electron dense outer layer, the exosporium, the classical electron-dense bumps shown in thick-exosporium spores, and hair-like projections, followed by the underlying spore coat, cortex, and core (Fig. 2A). Electron micrographs of EBB-or USD-treated spores show that both treatments were able to completely remove the spore coat and exosporium layers, leaving the spore-PG cortex exposed (Fig. 2A). Electron micrographs of Laemmli-treated spores reveal a thin layer of electron dense material surrounding the spore cortex, suggesting that some residual spore coat material remains attached to the cortex surface (Fig. 2A). Altogether, these results provide ultrastructural evidence that EBB and USD completely remove the spore surface layer after a single round of extraction, whereas Laemmli treatment leaves residual inner spore coat material.

**Figure 2.**
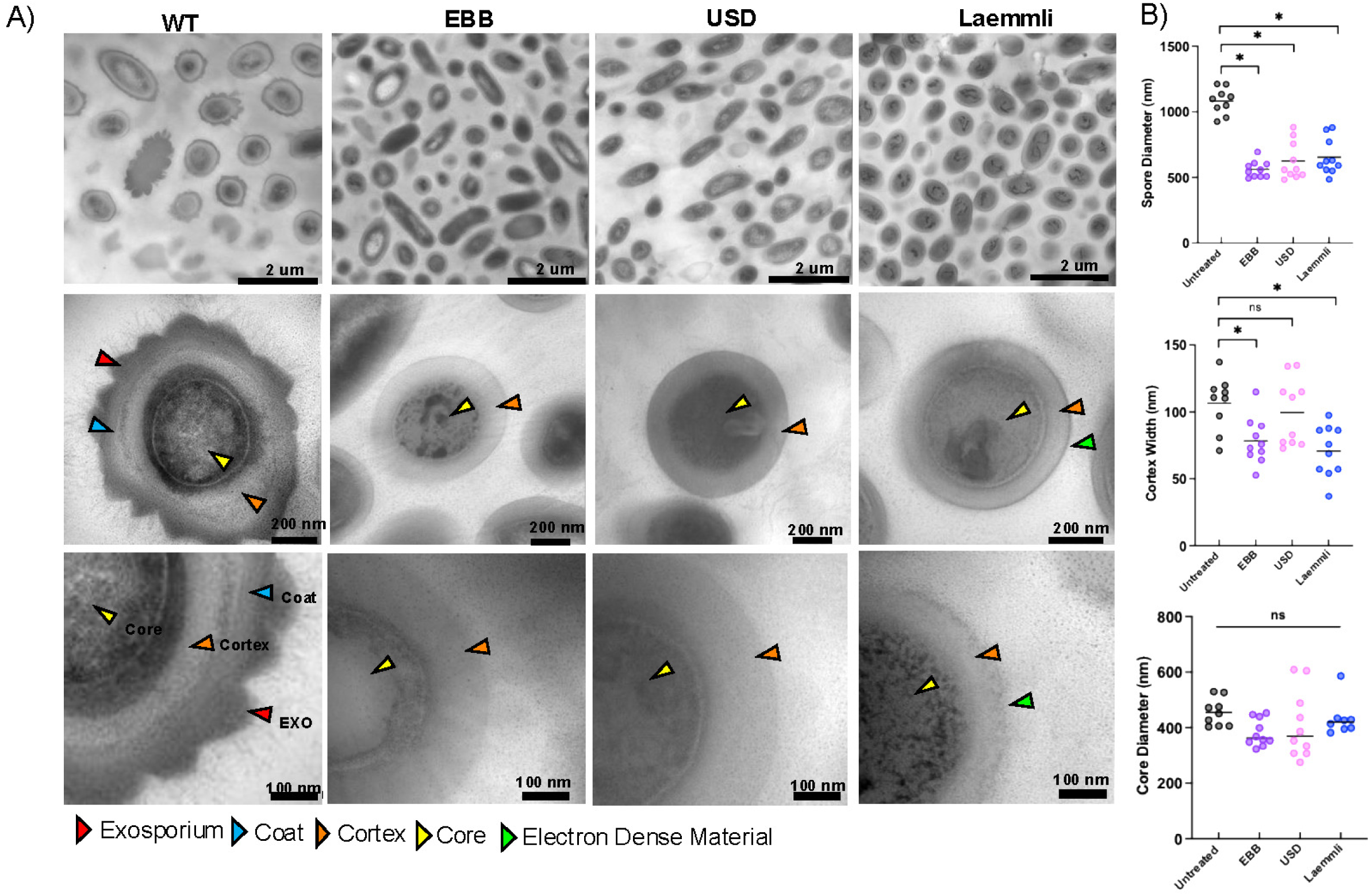
Transmission Electron Microscopy (TEM) Analysis of treated *C. difficile* spores. A) Representative images of wild type spores and spores treated with EBB, USD, or Laemmli buffers. Scale bars indicate 2 uM, 200 nm, and 100 nm respectively. Arrowheads point to: edge of exosporium (EXO,red), the coat (blue), cortex (orange), spore core (yellow), and residual electorn dense material (green). Measurements of spore diameter, cortex width, and core diameter were collected using ImageJ (B to D). 10 wild type spores and 10 of each treated spore were measured. Statistical significance was performed using a one-way ANOVA with Tukey’s multiple comparisons test. Significance is indicated as ns (not significant), * p < 0.05

To quantitatively assess the impact of these treatments on the ultrastructure of the spore layers, the spore diameter, cortex width, and core diameter were measured. The spore diameter significantly (p <0.05) decreased by ∼40% in spores treated with EBB, USD, and Laemmli when compared to untreated spores (Fig. 2B). *C. difficile* R20291 spores had an average diameter of ∼ 1,000 nm whereas on average Laemmli-, USD-, and EBB-treated spores were ∼ 600 nm diameter, attributed to the loss of the spore coat and exosporium layers (Fig. 2B). Comparison of the cortex width revealed several differences across treatments (Fig. 2B). A significant decrease was observed in cortex width between untreated spores and EBB (p<0.05) and Laemmli (p<0.05) treated spores respectively. On average, the cortex of untreated spores was ∼ 106 nm compared to ∼ 70 - 78 nm for EBB and Laemmli treated spores and 70 nm for Laemmli treated spores. No significant difference in cortex width between untreated spores (∼106 nm) and USD treated spores (∼100nm) was observed. Interestingly, despite the differences in cortex PG thickness, a slight but not significant decrease in spore core diameter was observed upon comparing USD, EBB, and Laemmli-treated spores to untreated spores (Fig. 2B). The decreased cortex width of EBB and Laemmli treated spores could be due to the boiling step during extraction, which leads to significant shrinkage in cortex width, this step is absent in USD-treated spores (Fig. 1). Boiling of spores after decoating releases dipicolinic acid (DPA) resulting in hydration and potentially release of proteins that might affect the cortex width ^17, 35^. No significant differences were observed in spore core diameter between untreated wild type spores and EBB, USD, or Laemmli treatments (Fig. 2B). Collectively, these results indicate that while all three treatments (i.e., EBB, USD, Laemmli) remove the spore coat and exosporium layer, those with a boiling step (i.e., Laemmli and EBB) affect the cortex width.

### Effect of EBB, USD, and Laemmli decoating in lysozyme-triggered cortex degradation

The spore coat layer is known to act as an impermeable barrier to molecules > 5 kDa, such as enzymes, including proteinase K and lysozyme ^32, 36, 37^. Moreover, the spore peptidoglycan cortex in decoated spores is susceptible to lysozyme degradation, artificially triggering spore germination and outgrowth ^32^. We reasoned that depending on the efficiency with which EBB, USD and Laemmli removes the coats, spores will have different levels of susceptibility to lysozyme-mediated cortex degradation. In the initial treatments, we observed differential impacts in the refractivity of *C. difficile* spores with spore cortex degradation seen in phase dark spores at 30 minutes incubation in treated samples while untreated spores remained phase bright (Fig 3A). While most of the EBB (64%%) and USD-treated spores (89%) remained phase bright after treatment (Fig. 3A), the majority (∼ 99%) of Laemmli-treated spores turned phase grey (Fig. 3B). Next, decoated spores where then treated with lysozyme (250 ug/mL) at 37°C for 30 min and 2 hours and observed under phase contrast microscopy (Fig. 3). Untreated control spores, ∼94% appeared phase bright before treatment and 93% remained phase bright after lysozyme treatment, indicating lysozyme was unable to degrade the PG cortex due to the spore coat remaining intact (Fig 3B). In contrast, ∼ 99% of EBB and USD treated spores became phase dark after lysozyme treatment, indicating effective coat removal allowing lysozyme triggered germination supported by TEM data (Fig. 3A,B). Interestingly, USD treated spores appeared to lyse leaving debris after 2 hr incubation with lysozyme. In the case of Laemmli decoated spores, subsequent lysozyme treatment resulted in 30% phase grey and 70% phase dark spores, indicating that Laemmli did not completely remove the spore coat from all the spores. Collectively, these observations demonstrate that EBB and USD treatment provides a homogenous population of decoated spores that can be utilized in downstream applications (e.g., studies of the PG cortex and/or spore core proteins).

**Figure 3.**
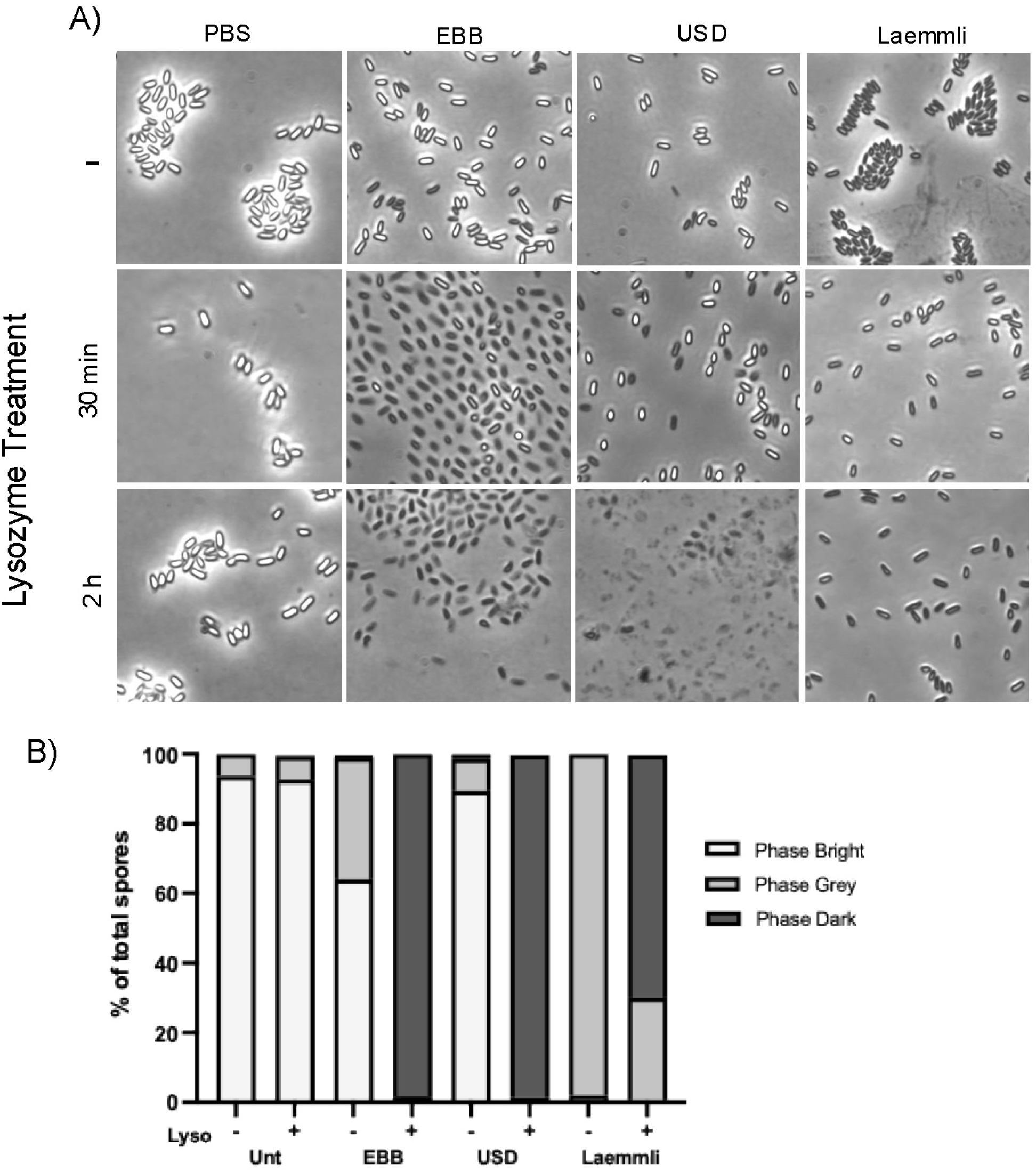
Effects of extraction treatment on spore coat integrity. A) Purified R20291 WT spores were treated with either EBB, USD, or Laemmli as previously described in Figure 1. Spores were treated with 250 ug/mL lysozyme and incubated for 30 min and 2 hours at 37°C. Lysozyme treated spores were imaged under phase contrast to assess effect of treatment on integrity of the spore coat and permeability to lysozyme. B) Spores for the 2 hour incubation were quantified using ImageJ cell counter function and categorized as phase bright, phase grey, or phase dark. Percentage in each category were graphed.

### Impact of EBB and USD-treatments in spore coat and exosporium removal

Having demonstrated that all three chemical treatments remove most of the spore surface layers, we thought to correlate the ultrastructure of spores treated with EBB, USD or Laemmli with the efficiency of extraction of spore protein markers by analyzing their protein-profile through SDS-PAGE. For this, spore extracts and decoated spores by SDS-PAGE electrophoresis (Fig. 1 and 4A,B). Notably, EBB spore extracts from a single EBB-treatment were able to extract the majority of spore proteins ∼90% within a single extraction (1^st^) (Fig. 1 and 4A, Fig. S1). Indeed, comparison of the remaining pellet of EBB-and USD/Laemmli-treated spores shows that EBB-treated spores have less residual spore coat / exosporium material attached (Fig. 1 and 4A). A second (2^nd^) and third (3^rd^) EBB treatment led to complete extraction of the spore coat and exosporium extracts and spore-pellets containing negligible remnants of spore coat and exosporium material (Fig. 1 and 4A, Fig. S1). In contrast, although a first USD-treatment yielded substantial protein extracts removing ∼74%, several high and low-molecular weight protein species remained in USD-spore pellets. A second (2^nd^) USD-treatment led to significant spore coat and exosporium extracts being removed, ∼18% with no additional detectable protein in a 3^rd^ extraction (Fig. 4B, Fig. S1). Notably, spore pellets from a second and third treatment with USD still retained detectable levels of unknown low molecular mass protein species (∼10 to 15 kDa) (Fig. 4B). Collectively, these results indicate that EBB is more effective than USD and Laemmli in the extraction of spore coat and exosporium proteins from *C. difficile* spores.

**Figure 4.**
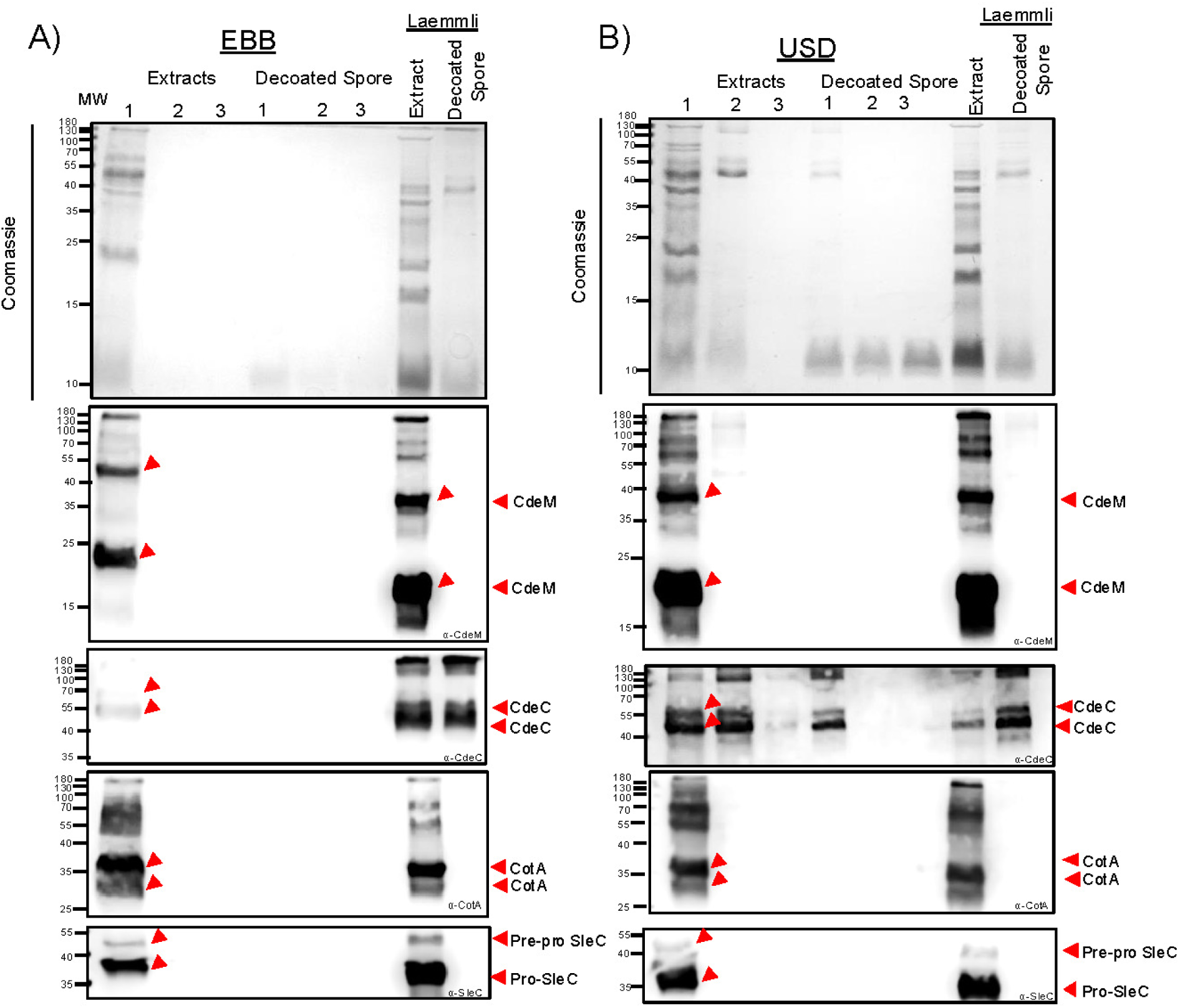
Extraction of *C. difficile* spore proteins using EBB, USD, or Laemmli. A) Purified R20291 WT spores were treated with consecutive extractions with EBB or USD and a single extraction with Laemmli buffer. Spores were treated as described in Methods and Figure 1. First (1^st^), second (2^nd^), and third (3^rd^) consecutive extractions and their respective decoated spores were resolved by 15% SDS-PAGE and gels were stained with Coommasie or subjected to immunoblotting with anti-CdeM, anti-CdeC, anti-CotA, and anti-SleC antibodies. B) anti-CdeM antibodies, arrows indicate immunoreactive bands at ∼70 kDa, 55 kDa, and 27 kDa respectively. anti-CdeC antibodies, arrows indicate immunoreactive bands indicating extraction of CdeC with bands at ∼130 kDa, 55 kDa, and 37 kDa, respectively. anti-CotA antibodies, arrows indicate extraction with bands ∼36 kDa. anti-SleC antibodies indicate immunoreactive bands at 55kDa pre-pro SleC and 37 kDa pro-SleC. Higher molecular weight bands likely represent oligomeric forms of the proteins.

### Impact of EBB and USD-treatments in spore surface markers

While the above protein profiles correspond to the electrophoretic separation of the total spore coat, exosporium, and cortex extracts, the extraction efficiency of each of these layers remains unclear. Therefore, we utilized antibodies against five proteins as markers. We considered a buffer to be efficient if it was able to remove the specific marker with a single extraction. For exosporium markers, we selected the cysteine-rich and morphogenetic proteins, CdeC and CdeM ^15, 17, 31^; where, CdeM is located on the outermost layer of the exosporium whereas CdeC is found within the exosporium with presumable interactions with the spore coat ^17, 38^. The essential morphogenic spore coat protein CotA was selected as a spore coat specific marker ^30, 39^. For the spore cortex marker, we selected the cortex-lytic enzyme SleC, known to be located in the cortex layer and essential for cortex degradation ^40^. Lastly, the germination protease GPR protein, necessary for the breakdown of small acid soluble proteins within the core was used a core marker ^19, 41^.

For this, aliquots of the 1^st^ EBB extraction were blotted against anti-CdeM shows immunoreactive bands at ∼23 kDa and ∼42 kDa likely corresponding to monomer and dimer forms of the protein (Fig. 4A). Higher oligomeric forms are also seen within the membrane (Fig. 4A). In the case of subsequent EBB extractions (2^nd^ and 3^rd^), there were no CdeM immunoreactive bands within the extracts or within the decoated spore pellets, indicating that the exosporium layer and CdeM protein was completely removed with a single treatment (Fig. 4A). The absence of exosporium was supported as seen through TEM (Fig. 2A). Extract of Laemmli-treated spores (1^st^) containing spore coat and exosporium extracts show the presence of ∼38 and ∼20 kDa immunoreactive species (Fig. 4A), while Laemmli spore-pellet completely lacked CdeM immunoreactive bands, it retained electron dense material in the spore surface as seen through TEM, likely not CdeM (Fig. 2A, Figure 4A). Of note, extraction with EBB led to differences in migration of the respective bands with EBB CdeM immunoreactive bands migrating slightly higher than those of the Laemmli extracted. Immunoblotting using anti-CdeM antibodies of 1^st^ USD spore extracts display immunoreactive bands at ∼ 38 kDa and ∼20 kDa and no subsequent CdeM immunoreactive band were observed in subsequent extractions 2^nd^ or 3^rd^ (Fig 4B). Moreover, no remnants of CdeM were detected in the USD-treated spore pellet (Fig. 4B). These results suggest that all three treatments efficiently remove the exosporium layer but differ in their levels of dissociation of the various CdeM oligomeric species present in the spore coat / exosporium extracts with USD and Laemmli dissociating CdeM as seen through the similar sizes of the immunoreactive bands whereas EBB leads to higher molecular weight immunoreactive bands of CdeM within the membrane.

Next, to explore whether CdeC, which is an exosporium protein suggested to be at the interface of the spore coat and the exosporium electron dense layer, was removed efficiently with a single treatment as as CdeM, immunoblots where analyzed using anti-CdeC antibodies ^12, 17^. Results suggest that a single extraction of EBB was sufficient to extract all immunodetectable CdeC of ∼44 kDa and 55 kDa, with no CdeC remnants in spore pellets (Fig. 4 A, B). However, in the case of USD-extracts, residual immunoreactive bands were detected in USD-decoated spores, which were fully removed after a second extraction with USD (Fig. 4B). These results suggest that EBB can completely extract CdeC with a single treatment compared to USD and Laemmli which retain CdeC material within the decoated spore following a single extraction. These results together suggest that EBB can extract and dissociate the exosporium and spore coat proteins as seen through CdeM and CdeC compared to USD and that Laemmli buffer, with the latter two requiring multiple rounds for complete removal of these proteins.

We also assessed the extraction of CotA as a spore coat protein marker^39, 42^. EBB extracts showed that a single extraction was sufficient to extract CotA with detectable immunoreactive bands detectable at ∼37 kDa and ∼34 kDa (Fig. 4A). No immunoreactive bands was observed after a second or third extraction in the exosporium/coat/cortex extracts or in the decoated spores (cortex/core). Interestingly, CotA was fully extracted with a single treatment with USD or Laemmli (Fig. 4B). No CotA immunoreactive bands were seen in USD extracts following a second and third extraction suggesting that most of the protein is removed with a single extraction (Fig. 4B).

A spore cortex protein, SleC, was used as a marker for extraction of cortex proteins^33^. During sporulation SleC is processed and appears as two immunoreactive bands in spore coat / exosporium extracts, with full length being ∼55 kDa and pro-SleC having a molecular weight of ∼37 kDa^40^. An initial EBB extraction displayed immunoreactive bands for both versions of SleC at the expected ∼55 kD and ∼37 kDa, while EBB-treated spore pellets did not have detectable SleC within the decoated spore (cortex/core fraction) (Fig 4A), indicating that SleC was completely extracted with a single treatment. In the case of USD, an initial USD-treatment extracted the majority of SleC with a single treatment similar to that seen in EBB and Laemmli treatments (Fig. 4B). In all three extractions methods SleC was completely removed with a single extraction with no detectable SleC in subsequent extractions or in decoated spore pellets. These observations indicate that a single with either treatment is sufficient to remove all of SleC from the spore cortex. Collectively, altogether, these observations suggest that EBB and USD are efficient in extracting exosporium, spore coat, and cortex markers and likely other proteins within these layers.

### Protein profile of EBB-and USD-treated spores following lysozyme treatment

The results showing that EBB and USD remove most of the spore coat and that these decoated spores are readily digested with lysozyme led us to speculate whether these steps could be utilized to lyse the spore core for electrophoresis of its contents. As observed in Figure 4 and Figure 5, three total extractions are sufficient to remove all residual spore surface markers. We repeated the three extractions with EBB and USD, observing similar levels of extraction of total protein and exosporium and coat markers CdeM, CdeC, CotA, or SleC in the first extraction (Fig. 5A,B, Supp Fig 1). Next, decoated spores were subjected to incubation with 2mg/mL lysozyme for 2 hours followed by electrophoresis and immunoblotting against all four exosporium, coat, and cortex markers as well as the spore core-specific marker, a germination protease (GPR). Significant protein contents were observable upon treatment of decoated spores with lysozyme in the SDS-PAGE. Immunoblotting against all four coat and exosporium markers yielded immunoreactive bands for CdeM and CdeC only in USD-, but not in EBB-decoated spores (Fig. 5A,B). It is noteworthy to clarify that the immunoreactive band observed in EBB and USD-decoated-lysozyme treated spores blotted with anti-CdeM is due to non-specific binding to lysozyme used during treatment. Notably, incubation of decoated spores with lysozyme was essential for detecting the spore core protein, GPR, as an immunoreactive band at the expected size of ∼37 kDa (Fig 5A,B). Altogether, these results demonstrate for spore core protein analyses, EBB allows extraction of spore core proteins, including GPR, in the absence of spore coat and exosporium.

**Figure 5.**
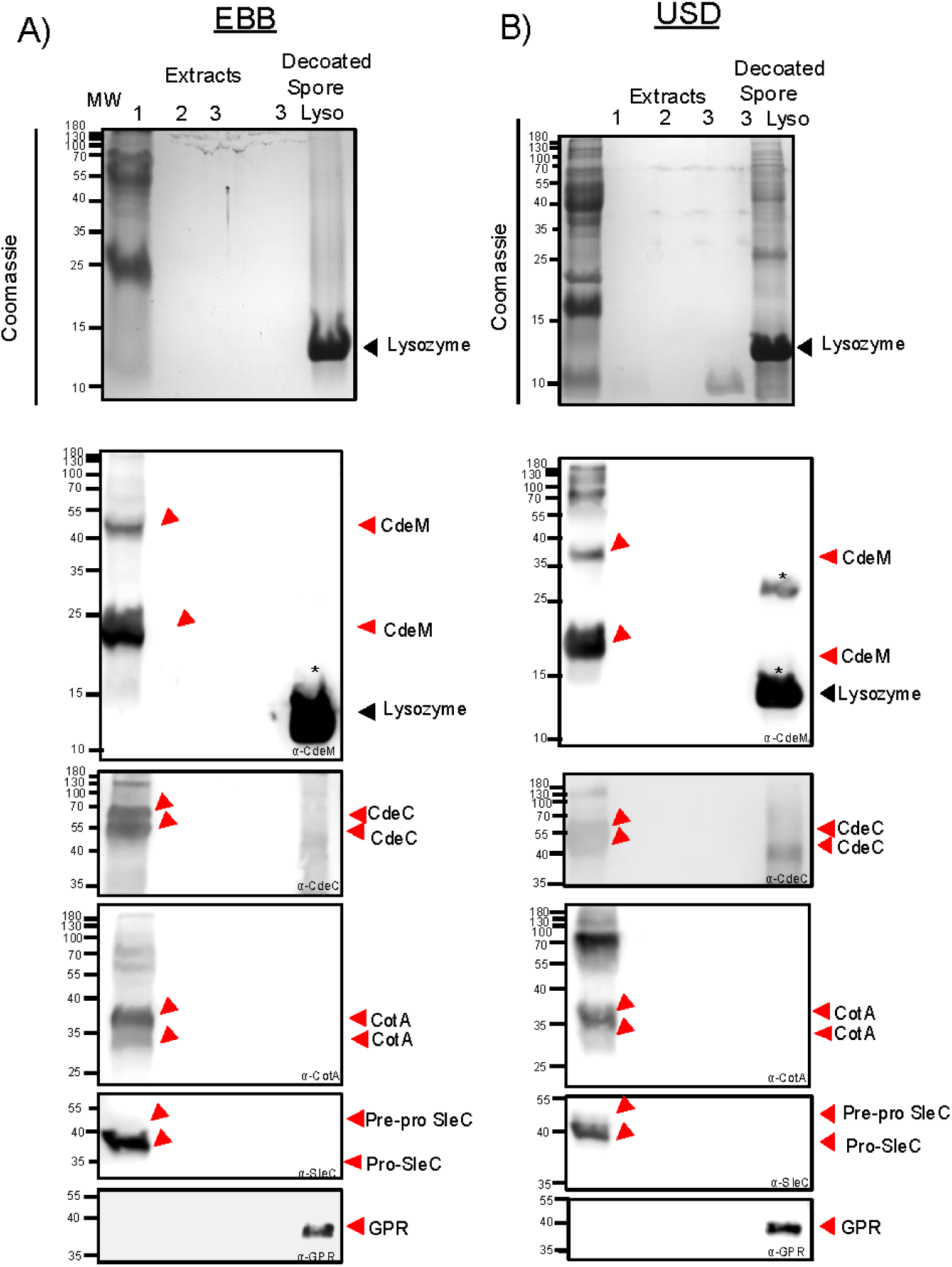
SDS-PAGE and Immunoblot analysis of lysozyme treated decoated *C. difficile* spores. Spore extracts from three subsequent rounds of extractions of EBB and USD treatments were resolved by 15% SDS-PAGE. In addition, fully decoated spores, following three rounds of EBB and USD. Decoated spores were incubated in 2 mg/mL lysozyme for 2 hours at 37°C to assess protein accessibility. Gels were stained with Coomassie and immunoblotted using anti-CdeM, anti-CdeC, anti-CotA, anti-SleC, and anti-GPR. Higher molecular weight bands represent oligomeric forms of the proteins. The ∼14 kDa molecular weight band corresponds to lysozyme is indicated. Non-specific binding of antibodies indicated by asterisk *.

## Conclusions

This work provides evidence that EBB-decoating treatment is more efficient than USD and Laemmli in removing the spore coat and exosporium layers. EBB also resulted in spore core protein extraction with no residual spore coat and exosporium. These techniques can be adapted to studies seeking fractionation of the different layers of the spore (i.e., exosporium, spore coat, spore peptidoglycan cortex, and spore core), immunoblotting, functional assays within *C. difficile* spores.

## Acknowledgments

This project was supported by awards 5R01AI116895 and 5R01AI172043 to J.A.S. and 5R01AI177842 to D.P-S. from the National Institute of Allergy and Infectious Diseases. The content is solely the responsibility of the authors and does not necessarily represent the official views of the NIAID. The funders had no role in study design, data collection and interpretation, or the decision to submit the work for publication.

